# Input-Specific Inhibitory Plasticity Improves Decision Accuracy Under Noise

**DOI:** 10.1101/2022.05.24.493332

**Authors:** Soomin C. Song, Bo Shen, Robert Machold, Bernardo Rudy, Paul W. Glimcher, Kenway Louie, Robert C. Froemke

## Abstract

Inhibitory interneurons regulate excitability, information flow, and plasticity in neural circuits. Inhibitory synapses are also plastic and can be modified by changes in experience or activity, often together with changes to excitatory synapses. However, given the diversity of inhibitory cell types within the cerebral cortex, it is unclear if plasticity is similar for various inhibitory inputs or what the functional significance of inhibitory plasticity might be. Here we examined spike-timing-dependent plasticity of inhibitory synapses from four major subtypes of GABAergic cells onto layer 2/3 pyramidal cells in mouse auditory cortex. The likelihood of inhibitory potentiation varied across cell types, with somatostatin-positive (SST+) interneuron inputs exhibiting the most potentiation on average. A network simulation of perceptual decision-making revealed that plasticity of SST+-like inputs provided robustness from higher input noise levels to maintain decision accuracy. Differential plasticity at specific inhibitory inputs therefore may be important for network function and sensory perception.

## Introduction

Synapses in the adult central nervous system are plastic and can be modified by changes in pre- and postsynaptic activity (Froemke, 2015; Magee and Grienberger, 2020; Malenka and Nicoll, 1999). Although most studies of long-term synaptic plasticity in the mammalian brain have focused on excitatory synapses, there is considerable evidence that GABAergic synapses are also plastic and express forms of inhibitory long-term potentiation (iLTP) and long-term depression (iLTD). Studies of inhibitory long-term plasticity in the rodent hippocampus (Horn and Nicoll, 2018; Huang et al., 2005; Saraga et al., 2008; Udakis et al., 2020; Woodin et al., 2003) and cortex (Chiu et al., 2018; D’amour and Froemke, 2015; Haas et al., 2006; Lagzi et al., 2021) have found that while sometimes inhibition can be adjusted in isolation from excitation, in general inhibitory plasticity seems to occur with activation or modification of excitatory synapses (Chiu et al., 2018; Field et al., 2020). Coordinating plasticity across different synapse types could be important for regulating a variety of single-cell integrative properties including excitatory-inhibitory balance, spike timing, and dynamic range of spike generation (Chiu et al., 2019; Hattori et al., 2017; Isaacson and Scanziani, 2011; Maffei et al., 2017; Vogels et al., 2013).

Studies of inhibitory plasticity in the cerebral cortex historically have been challenging in part because of the diversity of GABAergic interneuron subtypes (Gouwens et al., 2020; Huang and Paul, 2019; Tremblay et al., 2016). This diversity endows cortical circuits with powerful computational flexibility. However, functionally-distinct inhibitory cells or cell types might then be expected to also have different plasticity rules governing the induction and expression of iLTP and iLTD. Distinct learning rules for inhibitory plasticity might also in turn contribute to the different functional roles assigned to specific inhibitory inputs or sub-circuits, e.g., with varying requirements for changes in activity levels over a range of timescales, and dependent on variations in synaptic connectivity and short-term plasticity (Froemke et al., 2010a). Conventionally there have been three major classes of cortical interneurons (Callaway 2016; DeFelipe et al., 2013; Fishell and Rudy, 2011; Kuchibhotla et al., 2017), defined by expression of the molecular markers parvalbumin (PV), somatostatin (SST), and vasoactive internal peptide (VIP). More recent work has begun to describe other cortical interneuron types largely but not exclusively located in superficial layers, such as cells selectively expressing neuron-derived neurotrophic factor (NDNF) among others (Abs et al., 2018; Cohen-Kashi Malina et al., 2021; Schuman et al., 2019; Tasic et al., 2016). These subtypes have different spontaneous and evoked patterns of action potential firing resulting from various complements of ion channels and receptors, form synapses onto target pyramidal cells in specific dendritic domains, and are recruited more or less in phase with excitatory cells (Dorrn et al., 2010; Higley and Contreras, 2006; Wehr and Zador, 2003). As these cellular properties influence the induction and expression of plasticity of excitatory synapses (Froemke et al., 2010b), it is likely that inhibitory plasticity would also then vary by synapse type and thus cell type. In agreement with this, recent work on long-term inhibitory plasticity in frontal cortex (Chiu et al. 2018; Lagzi et al., 2021) and hippocampus (Udakis et al. 2020) found that similar induction procedures led to different results at synapses formed by PV+ compared to SST+ interneurons; in both cases, iLTP was more readily induced at SST+ inputs onto excitatory cells.

Previously we showed that pairing pre- and postsynaptic action potentials led to spike-timing-dependent plasticity (STDP) of excitatory and inhibitory postsynaptic potentials (E/IPSCs) elicited by electrical extracellular stimulation in slices of mouse auditory cortex (D’amour and Froemke, 2015; Field et al., 2020). However, these electrically-evoked IPSCs likely constituted a mixed population of inputs from the main interneuron types. Here we aimed to compare the efficacy of pre- and postsynaptic spike pairing for induction of spike-timing-dependent iLTP in PV+, SST+, VIP+, and NDNF+ inputs in superficial layers of mouse auditory cortex. Additionally, these inputs have different postsynaptic targets, with SST+/PV+ inputs providing feedback/feedforward inhibition to local excitatory pyramidal cells, and VIP+ cells disinhibiting excitatory neurons via inhibition onto other interneurons (Karnani et al., 2016; Kepecs and Fishell, 2014; Lee et al., 2013; Pfeffer et al., 2013; Zhang et al., 2014). As it is not necessarily obvious how changes in excitatory and inhibitory inputs act together to affect overall network performance, we also used a circuit model of cortical connectivity simulating a decision-making process based on noisy inputs to ask about the functional consequences of potential plasticity at these various inputs.

## Results

### Spike Pairing Differentially Induces LTP Across Four Types of Inhibitory Inputs

Our primary experimental goal was to determine the potential for iLTP induction at the various classes of inhibitory inputs onto cortical excitatory neurons. We made whole-cell recordings from layer 2/3 pyramidal neurons in slices of auditory cortex of adult mice. An extracellular stimulation electrode was placed laterally from the recording region in layer 2/3 and used to evoke mixed synaptic responses onto the recorded cell (**Figure 1A**). Most recordings were made from transgenic mice virally expressing channelrhodopsin-2 specifically in cortical interneurons of a specific subtype. For each recording, we monitored IPSC amplitude in voltage-clamp mode (at a holding potential of −40 mV), recording either electrically-evoked composite IPSCs and/or optically-evoked IPSCs by optogenetic stimulation with blue light through the microscope objective. After measuring baseline events for 5-10 minutes, recordings were switched to current-clamp to pair electrically-evoked inputs with single postsynaptic spikes a total of 100 times (Bi and Poo, 1998; D’amour and Froemke, 2015; Feldman, 2000). After pairing, recordings were returned to voltage-clamp mode and electrically/optically-evoked IPSCs were monitored for the duration of the recording (**Figure 1B**).

**Figure 1.**
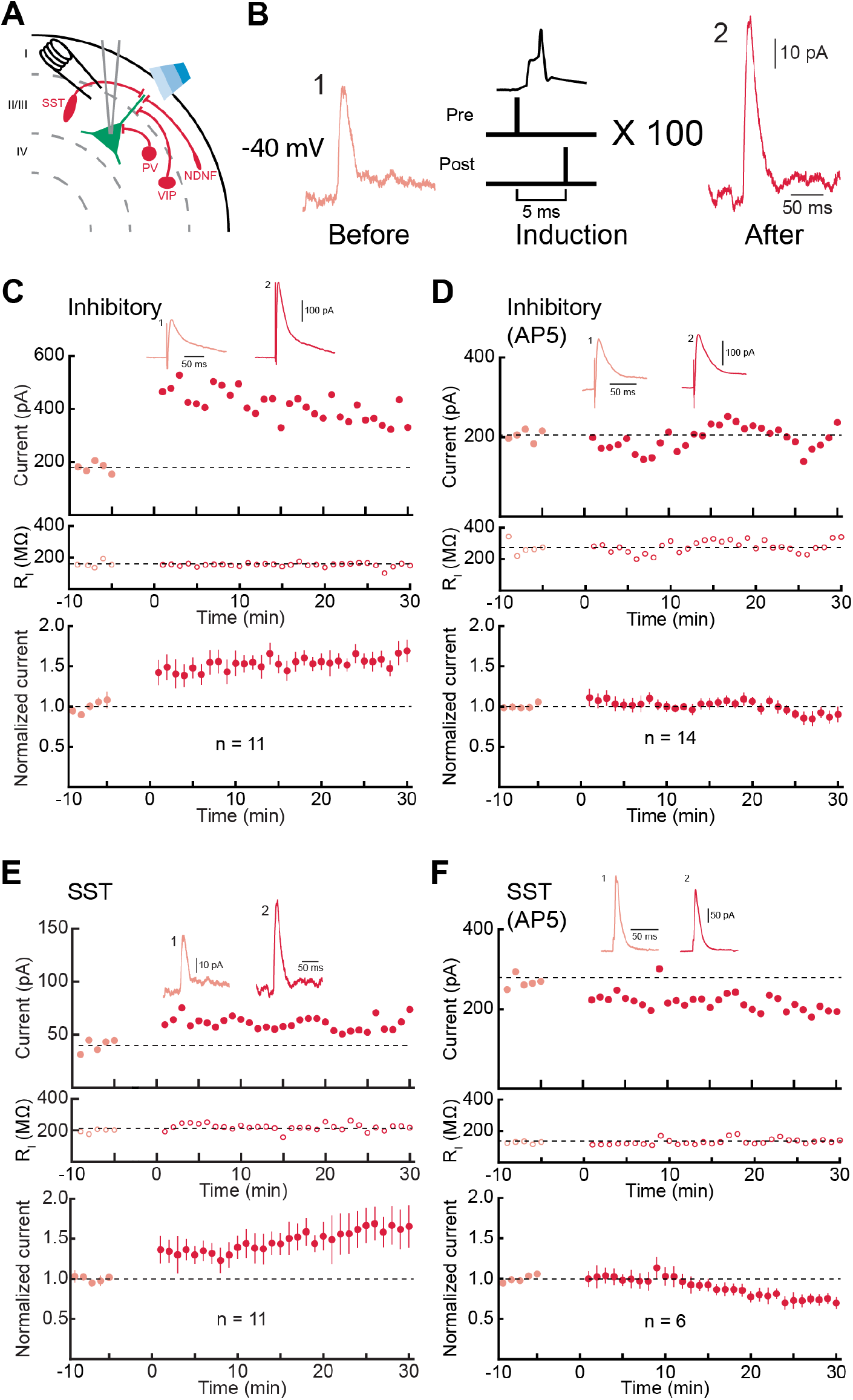
SST+ inhibitory inputs are reliably potentiated by spike pairing. (**A**) Whole-cell recordings from mouse auditory cortical layer 2/3 pyramidal cells in slices using extracellular electrical stimulation and optogenetics to elicit synaptic responses. (**B**) Protocol for attempting to induce iLTP with pre→post spike pairing. Before and after pairing, IPSCs were recorded in voltage-clamp mode at −40 mV; inputs were paired with postsynaptic depolarization to trigger single action potentials in current-clamp mode for 100 trials. (**C**) Potentiation of electrically-evoked IPSCs by spike pairing. Top, example recording from wild-type mouse auditory cortex with IPSCs evoked by extracellular electrical stimulation. Dashed line, pre-pairing mean. Insets show example IPSCs before pairing (1) and 11-20 minutes after pairing (2). Middle, input resistance (R_i_) for this cell. Bottom, summary of IPSC amplitude before and after spike pairing (n=11 cells). Error bars, SEM. (**D**) NMDA receptor antagonist AP5 (50 μM) prevented iLTP induction for electrically-evoked IPSCs. Top and middle, example cell IPSCs and input resistance. Bottom, summary. (**E**) iLTP at optically-elicited SST IPSCs in slices from SST-Cre mouse auditory cortex with channelrhodopsin-2 expressed in SST+ cortical neurons. Top and middle, example cell IPSCs and input resistance. Bottom, summary. (**F**) AP5 (50 μM) also prevented iLTP of SST IPSCs. Top and middle, example cell IPSCs and input resistance. Bottom, summary.

We found that spike pairing induced a robust LTP of electrically-evoked IPSCs (**Figure 1C**), similar to the inhibitory STDP previously described onto layer 5 pyramidal cells (D’amour and Froemke, 2015). For the cell shown at the top of **Figure 1C**, pre→post spike pairing led to an increase of IPSC amplitude of 237.7% from baseline as measured at 11-20 minutes after pairing. Over 11 cells, the average iLTP was 163.2±10.3% of the baseline synaptic strength (**Figure 1C**, bottom; p<0.001, Student’s paired two-tailed t-test). Similar to inhibitory plasticity in auditory cortex layer 5 and hippocampal CA1 neurons (D’amour and Froemke, 2015; Huang et al. 2005), iLTP was prevented if NMDA receptors were blocked with bath application of 50 μM AP5 (**Figure 1D**; iLTP: 105.2±4.8% of baseline, n=14, p=0.43). Therefore, coordinated activation of excitatory and inhibitory inputs together with postsynaptic spiking acts to enhance the strengths of those co-activated inhibitory synapses.

In brain slices prepared from various GABAergic interneuron Cre lines, we then asked how spike pairing modified inhibitory subtype-specific IPSCs. STDP was induced in the same way (pairing postsynaptic spiking with electrically-evoked synaptic inputs), but before and after pairing we used optical stimulation to monitor SST-IPSCs, PV-IPSCs, VIP-IPSCs, or NDNF-IPSCs. We found that SST-IPSCs were highly plastic (**Figure 1E**; SST iLTP: 155.2±15.4% of baseline, n=11, p=0.006), modified to much the same degree as composite electrically-evoked IPSCs. This included the sensitivity to NMDA receptor blockade, as 50 μM AP5 also prevented spike pairing from inducing SST iLTP (**Figure 1F**; iLTP: 95.3±7.2% of baseline, n=6, p=0.47).

Other classes of inhibitory inputs showed a range of responses to pre→post spike pairing. The magnitude of changes at these other synapses was much smaller than iLTP at SST inputs. PV-IPSCs and NDNF-IPSCs could be somewhat potentiated by spike pairing (**Figure 2A,B**; PV iLTP: 126.4±6.8%, n=9, p=0.005; NDNF iLTP: 113.4±5.2%, n=9, p=0.045). In contrast, VIP-IPSCs were not significantly affected across the population by spike pairing in this manner (**Figure 2C**; VIP iLTP: 115.5±16.3%, n=7, p=0.7). Overall, all 11/11 cells recorded with electrically-evoked IPSCs monitored showed significant iLTP of the evoked events; this dropped to 3/14 cells showing significant iLTP when NMDA receptors were blocked. Similarly, 10/11 pyramidal cells with SST-IPSCs monitored had significant iLTP, but significant iLTP was observed in only 4/9 cells with PV-IPSCs monitored, 4/9 cells for NDNF-IPSCs, and 2/7 cells for VIP-IPSCs (**Figure 2D**). These data show that after repetitive pairing of pre- and postsynaptic activity, four of the major classes of inhibitory input onto layer 2/3 pyramidal cells had varying tendencies to be reliably potentiated.

**Figure 2.**
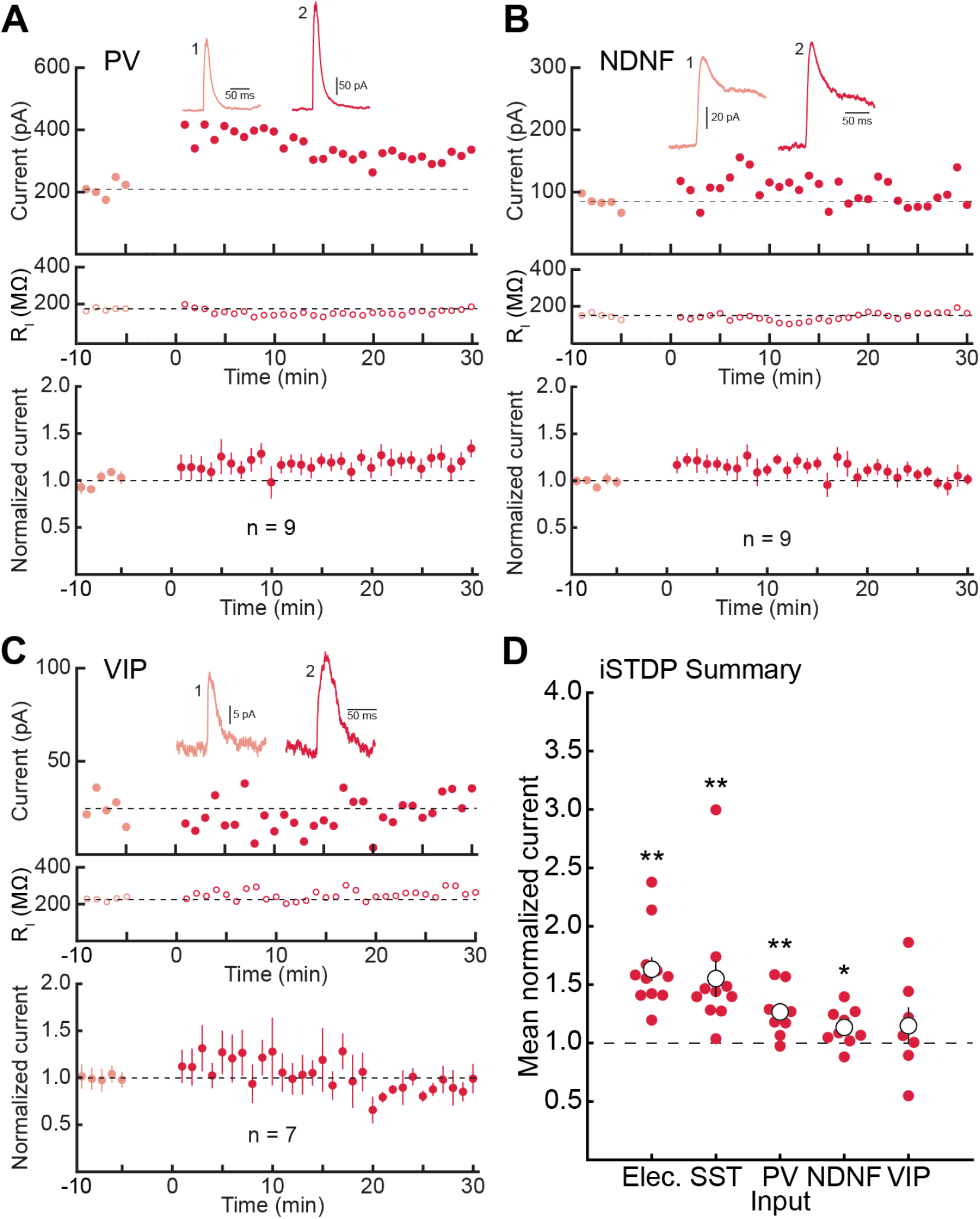
Pairing induces iLTP of PV+ and NDNF+ but not VIP+ inputs to excitatory neurons. (**A**) iLTP at optically-elicited PV IPSCs. Top and middle, example cell IPSCs and input resistance. Bottom, summary. (**B**) iLTP at optically-elicited NDNF IPSCs. (**C**) Lack of iLTP at optically-elicited VIP IPSCs. (**D**) Summary of iLTP induction attempted for different electrically- or optically-evoked IPSCs. Red symbols, individual recordings; open circles, mean amplitude change post-induction. **, p<0.01; *, p<0.05. Error bars, SEM.

### Simulation of Perceptual Decision Making

We then used a modeling approach based on Shen et al. (2022) to examine functional consequences of differential plasticity at various types of inputs. We tested the behavioral implications of iLTP using a biologically-based circuit model of decision-making (**Figure 3A**). The model consisted of three types of units which simulate pools of cortical neurons: excitatory (R units), inhibitory (G units), and disinhibitory (D units). Excitatory R units represent cortical pyramidal neurons, receiving both the external inputs and providing the final output. Inhibitory G units represent SST+ inhibitory neurons (and to some degree, PV+ interneurons) that provide feedback gain control to local pyramidal neurons, whereas disinhibitory D units represent cortical VIP+ neurons that usually inhibit SST+/PV+ neurons and therefore disinhibit the pyramidal neuron population (Fu et al., 2014; Karnani et al., 2016; Kepecs & Fishell, 2014; Lee et al., 2013; Pfeffer et al., 2013; Pi et al., 2013; Tremblay et al., 2016).

**Figure 3.**
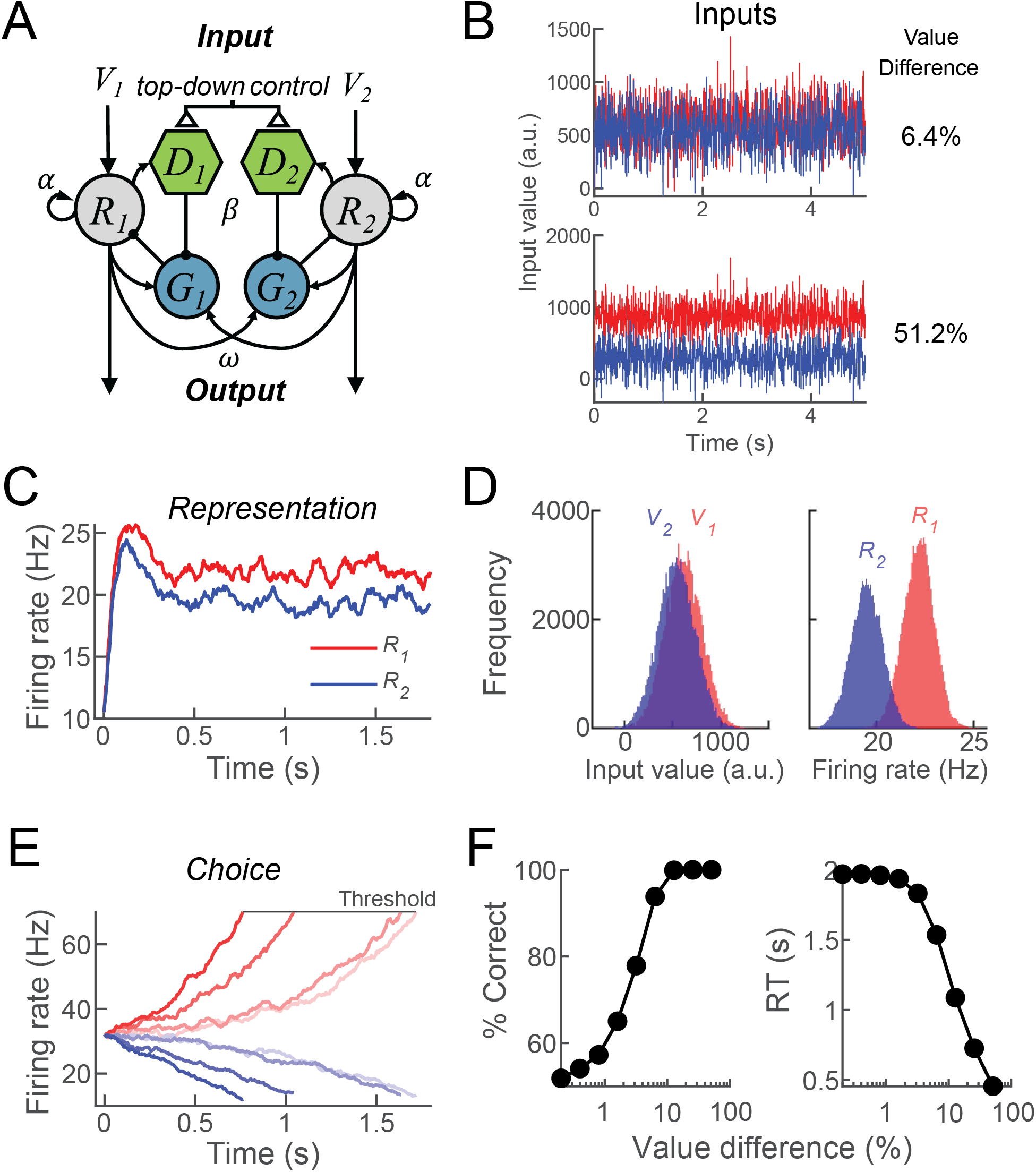
Generalized cortical circuit model of inhibitory and disinhibitory processing. (**A**) The circuit model consists of two pools of units, each receiving a noisy input V onto an excitatory R unit. Each R unit also has recurrent excitation α, receives inhibitory input G from the same pool and sends excitatory inputs to the G units of both pools. Each D unit receives excitatory input from the local R unit with the R-D connection gated by top-down control during the ‘choice’ but not the ‘representation’ mode of the model, and sends an inhibitory input to the G unit of the same pool. (**B**) Examples of input values over time for V1 (red) and V2 (blue) for two different value differences. (**C**) Example simulation run in representational mode when input V1 is slightly larger on average than V2. Shown are the firing rates over time of the two R units; diverging and reaching steady state with accurate representation of the relative input strengths. (**D**) Distribution of input values V1 and V2 (left) and steady-state firing rates of R1 and R2 (right) during the simulation in **C**. (**E**) Four example simulation runs in choice mode when V1 and V2 have varying value difference (higher value difference, darker red and blue traces; lower value difference, lighter red and blue traces) (**F**) Simulated psychometric choice behavior (left) and chronometric reaction time (right) as functions of V1 vs V2 value difference.

The goal of the model was to receive a noisy sensory input and classify the input as belonging to one of two categories. Each run of the simulations proceeded in one of two modes: a ‘sensory representation’ mode with the disinhibitory D units not activated, and a ‘behavioral choice’ mode with D units active. This modeling approach was designed to emulate the task structure of two-alternative forced-choice tasks used to test input discriminability and reaction times. During the simulation, each of the two excitatory R units received a continuous noisy input about the stimulus value of one of two choices V1 and V2 (**Figure 3B**). The two R units each had self-excitation (α) in addition to the excitatory input V1 or V2, and received inhibitory input from one of the G units. Each G unit received excitatory projections from both R units, enabling lateral inhibition between the two R units. Each disinhibitory D unit received excitatory input from one of the R units, with this input inactivated (set to 0) during the ‘representation’ mode of the model. This simulated the gating of VIP+ disinhibition via top-down control or an external modulatory input that allowed for non-zero excitatory input during the ‘choice’ mode of the model. The D units in turn inhibited the inhibitory G units, therefore disinhibiting the local R unit and driving competition between the Rs to result in a winner-take-all choice process between the ongoing noisy external inputs V1 and V2 (**Figure 3A**). The decision process was challenged by varying the value difference (defined as: (|V1-V2|) / (V1+V2)) between the V1 and V2 inputs, making them more or less similar as a function of the noise level (**Figure 3B**).

Example simulations are shown both for the representational mode without disinhibition (**Figure 3C,D**) and for the choice decision-making mode with active disinhibition (**Figure 3E,F**). During the representational mode, the dynamics of R1 and R2 firing rates show transient peaks before settling to an equilibrium state that accurately represented the relative value of the two noisy inputs. Even when the two inputs are highly overlapping, the representations rapidly diverge (**Figure 3C**) and the firing rates of R1 and R2 units are well separated (**Figure 3D**).

During the choice mode, active disinhibition leads to emergence of a winner-take-all dynamic in the model between the two pools of units (R1,G1,D1 vs R2,G2,D2). Over the course of the simulation, the pool of units receiving stronger inputs tended to ramp up and reach a decision threshold while the activity of the other pool was suppressed (**Figure 3E**). This means that after a period of deliberation, input integration turned into a categorical choice of stronger input when the neural activity of one of the R units hit an arbitrary decision boundary. Over multiple simulations we quantified two properties of the model, choice accuracy (the proportion of trials in which the R unit receiving the larger input hit the decision boundary) and reaction times (number of time steps until the decision threshold was reached). Consistent with standard empirical findings in decision-making tasks (Roitman and Shadlen, 2002; Barack and Gold 2016), choice accuracy increased and reaction time decreased as functions of value difference, i.e., the contrast between V1 and V2 (**Figure 3F**).

### Inhibitory Potentiation Improves Decision-Making in Noise

Finally, we asked if inhibitory plasticity might improve decision making in the model. We hypothesized that iLTP from the G units (simulating SST+/PV+ interneurons) onto the R units (simulating pyramidal neurons) in the network might enhance the effective signal-to-noise ratios, improving overall integration and choice between V1 from V2 especially for higher noise levels. We tested this hypothesis by examining model performance under different connection weights from the G to the R units in the network. First, we examined how increasing inhibition (simulating iLTP) from G onto R affects the representational mode with disinhibition inactive. As expected, iLTP decreased the firing rates of R1 and R2 (**Figure 4A**, filled curves) compared to baseline (**Figure 4A**, dashed curves). Choice accuracy increased with greater amounts of iLTP (**Figure 4B**). By examining the reaction times of the model, we found that the improvement of choice accuracy emerged from the prolonged deliberation time on choice of V1 vs V2 (**Figure 4C**). Improvement of choice accuracy was more prominent when the input noise on the V1 and V2 signals was higher (**Figure 4D**).

**Figure 4.**
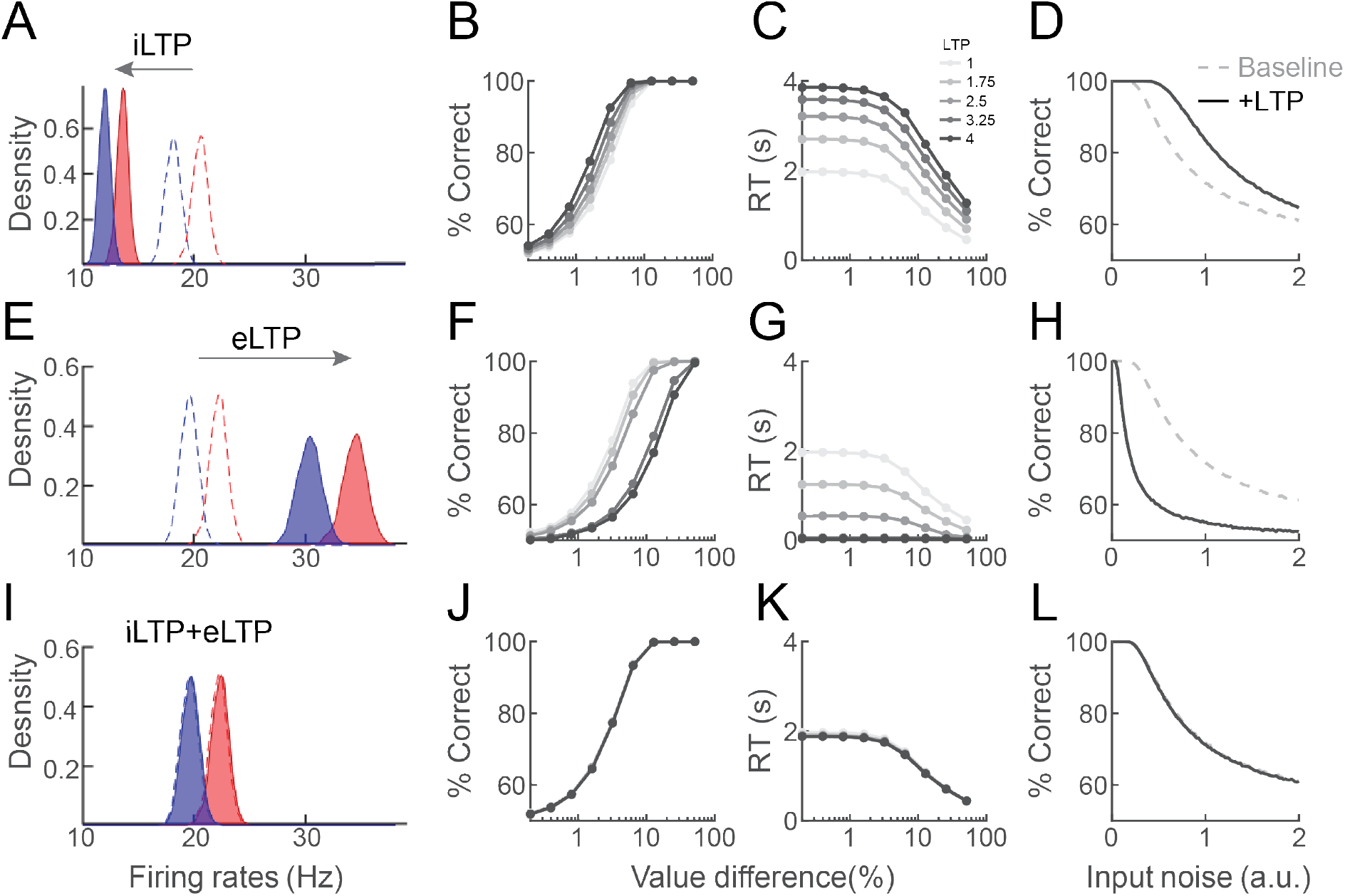
iLTP improves choice accuracy in noise. (**A**) iLTP was simulated by increasing the connection strength from the G onto the R units. Shown are the firing rate distributions for one simulation in representation mode, depicting baseline firing rates of R1 (red) and R2 (blue) before iLTP (dashed lines) and after iLTP (filled distributions). iLTP alone decreased R unit firing rates. (**B**) Performance of the model in choice mode (in terms of classification % correct) as a function of value difference between the V1 and V2 inputs. Larger contrast between the V1 and V2 values led to better classification performance. For intermediate levels of value difference, iLTP of different magnitudes (increasing from gray to black lines) improved performance. (**C**) Larger amounts of iLTP increased the reaction time (RT) until the model reached choice decision threshold. (**D**) Choice accuracy as a function of input noise level, under baseline conditions (dashed line) and after iLTP (solid line). Improved choice accuracy with iLTP is more prominent at higher noise levels. (**E**) As in **A**, but for eLTP (simulated by increasing the V-to-R and α connection strengths). eLTP alone increased R unit firing rates. (**F**) As in **B**, but for eLTP. eLTP of different magnitudes (from gray to black lines) impaired model choice accuracy. (**G**) As in **C**, but for eLTP. Larger amounts of eLTP decreased RT. (**H**) As in **D**, but for eLTP. Impaired choice accuracy with eLTP at all noise levels. (**I**) As in **A**, but for iLTP together with eLTP (iLTP+eLTP). Firing rates of the R units were left intact after adjusting synaptic strengths. (**J**) As in **B**, but for iLTP+eLTP. Choice accuracy was preserved at baseline levels when iLTP and eLTP are balanced. (**K**) As in **C**, but for iLTP+eLTP. RT was preserved at baseline levels. (**L**) As in **D**, but for iLTP+eLTP. Across noise levels, model choice accuracy was preserved.

Enhanced inhibition in the network model might indirectly result from increased excitation of the R units as well. Therefore, we examined the consequences of strengthening excitation instead of inhibition, i.e., simulating the effects of excitatory LTP (eLTP), either without iLTP (**Figure 4E-H**) or combined with iLTP (**Figure 4I-L**). For eLTP, we increased the connection strengths both of the input signals (V1 and V2) onto the R units, as well as recurrent self-excitation (α). eLTP led to higher R unit firing rates during the representation mode (**Figure 4E**), but decreased accuracy during the choice mode (**Figure 4F**). Decreased choice accuracy after eLTP resulted from a shorter reaction time to reach decision threshold (**Figure 4G**), and was exacerbated particularly at moderate-to-higher noise levels of the input (**Figure 4H**). However, if eLTP was combined with iLTP, baseline firing rate distributions were preserved (**Figure 4I**), and choice accuracy was restored to the original levels before eLTP (**Figures 4J-L**). Thus, iLTP can enhance the deliberative process of decision making particularly for noisy inputs. When coordinated with eLTP processes presumably critical for learning and memory, we suggest that iLTP at specific synapses can help correct for performance impairments produced by eLTP and increases in overall excitability.

## Discussion

Inhibitory plasticity endows neural circuits with a degree of flexibility surpassing that provided only by conventional excitatory plasticity (Froemke, 2015; Haas et al., 2006; Lagzi et al., 2021; Luz and Shamir, 2012; Vogels et al. 2013). Here we documented how pre- and postsynaptic spike pairing led to differing degrees of iLTP at four major classes of inhibitory inputs onto layer 2/3 pyramidal cells in mouse auditory cortex. We found that SST+ inputs were most plastic, suggesting that potentiation of these inputs is predominantly responsible for changes to mixed populations of electrically-evoked IPSCs. An alternative hypothesis is that electrical extracellular stimulation mainly recruits SST+ cells, at least in brain slices of mouse auditory cortex. Spike pairing also induced iLTP at PV+ and NDNF+ inputs to a much lesser extent, while VIP+ inputs were not significantly affected. This observation of heterogeneous iLTP across cell types is similar to a recent study in mouse prefrontal cortex (Chiu et al., 2018), with inhibitory plasticity induced by NMDA application mainly at SST+ inputs. In principle, different cell and synapse types should have distinct learning rules to help ensure the functional specialization of specific interneuron classes. This almost certainly must be the case due at least to differences in the connectivity patterns between cell types, which should impact how pre- and postsynaptic events are integrated in the pyramidal neuron dendrites (Froemke et al. 2005; Froemke et al. 2010b). Indeed, if the mechanistic requirements for long-term plasticity were identical across all inhibitory inputs, this might dissolve any functional specificity and ensure a homogeneous distribution of inhibitory responses irrespective of molecularly-defined cell type identity.

We designed a straightforward network model of cortical connectivity and decision making to assess potential functional significance of iLTP. Inhibitory potentiation was modeled as an increase in connection strength from the G to the R nodes, simulating iLTP of SST+ and possibly PV+ feedforward inputs onto local pyramidal neurons. We found that this led to longer ‘reaction times’ for either R node to reach the decision threshold, naturally improving choice accuracy especially for higher noise levels. This essentially rendered the model less sensitive to the effects of sudden high-amplitude input transients. This is just one of several potential functions of inhibitory potentiation for regulating neuronal and network excitability, and in addition there are presumably other conditions in which iLTD (instead of iLTP) might be functionally advantageous. For example, in dynamic environments, false positives at the expense of slower reaction time and accuracy might be important for decision making (Glennon et al., 2019). Correspondingly, SST+ and PV+ interneurons throughout the cortex and hippocampus are directly regulated by a number of neuromodulators including acetylcholine, noradrenaline, and oxytocin among others (Krugliokv and Rudy, 2008; Kuchibhotla et al., 2017; Lee et al., 2010; Marlin et al., 2015; Muñoz and Rudy, 2014; Nakajima et al., 2014; Salgado et al., 2010). These modulators can act to bi-directionally control interneuron synaptic and spiking activity over a range of timescales, which would be analogous to the change in gain of inhibition in the model, as well as enabling forms of excitatory and inhibitory long-term plasticity themselves (Froemke et al., 2013; Froemke, 2015; Kirkwood et al., 1999; Martins and Froemke, 2015; Seol et al., 2007).

We focused on the specific contributions of inhibitory plasticity, as much less has been known about iLTP compared to the large literature on the mechanisms and potential functional relevance of excitatory long-term synaptic plasticity. This includes several reports of STDP and eLTP at recurrent or lateral connections in layer 2/3 of the rodent cortex (Froemke and Dan, 2002; Karmarkar et al., 2002). For these reasons we did not further examine the functional contributions of eLTP or other forms of excitatory plasticity for network function beyond input integration in our model. There is also considerable evidence that if eLTP is the sole unchecked mechanism for changing network weights, the overall increase in net excitability can dominate network performance and impair overall function (Davis, 2006; Wu et al., 2020; Zenke et al., 2016). While other mechanisms such as lagged synaptic scaling or other more global homeostatic mechanisms can in theory compensate for these excitability changes, some of these mechanisms are too slow or non-specific to appropriately maintain both network excitability and performance. We have recently described how homosynaptic and heterosynaptic inhibitory plasticity can provide rapid and input-specific regulation of excitatory-inhibitory balance (D’amour and Froemke, 2015; Field et al., 2020). These previous studies neglected the important contributions and distinctions of separate inhibitory subtypes, which motivated the experiments here. In agreement with past work from our labs and others (Haas et al., 2006; Lagzi et al. 2021; Luz and Shamir, 2012; Vogels et al. 2011; Vogels et al., 2013), inhibitory plasticity can be computationally advantageous, but may be most effective when coordinated with excitatory changes to prevent reductions in firing rate and dynamic range which also might have negative consequences if left uncorrected. It is important to note that many studies have found that inhibitory long-term plasticity depends on NMDA receptor activation (i.e., excitatory input signaling), helping to ensure that mechanisms of excitatory and inhibitory modifications are engaged together. Precisely how NMDA receptor signaling leads to appropriate adjustment of co-activated GABAergic synapses remains a major outstanding question in the field.

Our model was designed to operate in two modes, specifically to help determine the consequences of input-specific plasticity of the G units onto the R units. The two modes were switched from representational to choice by including a disinhibitory mechanism (the D unit projection onto the G units), simulating the activity of VIP+ neurons in cortical networks. Gated disinhibition provides a biologically-plausible mechanism for top-down control, supported by experimental evidence of long-range inputs and context-dependent neuromodulation of VIP+ interneurons (Fu et al., 2014; Kamigaki, 2019; Lee et al., 2013; Pi et al., 2013; Pinto and Dan, 2015; Zhang et al., 2014), which can powerfully promote plasticity of other co-activated synapses (Canto-Bustos et al., 2022). We found that VIP+ inputs onto layer 2/3 pyramidal cells were not reliably affected by spike pairing. Thus we did not include a connection from the D unit back to the R unit, as this input was not modified under these conditions and we primarily aimed to determine the consequences of changing the strengths of the G units. Our data do not imply that VIP+ cells and synapses are absolutely implastic; there are potentially other induction procedures or mechanisms that might lead to iLTP or iLTD of these connections as well. Spike pairing is one protocol among many for synaptic plasticity induction, and it is well established that changing the timing and/or amount of pre- and postsynaptic activity can modify synapses otherwise insensitive to more innocuous procedures (Froemke et al., 2010a; Gomez et al., 2014). It is also possible that an important site of plasticity for VIP+ inhibitory inputs is onto the direct SST+ interneuron postsynaptic targets responsible for disinhibition. Our studies provide a step towards understanding the complete logic of inhibitory synaptic plasticity rules, how these changes are coordinated with sets of excitatory inputs, and the net consequences for network function that emerge from these dynamics.

## Acknowledgements

We thank D. Lin and R.W. Tsien for comments, discussions, and technical assistance. This work was funded by grants from the NINDS (NS074972) to B.R. and R.C.F., and NIDCD (DC012557), NICHD (HD088411), and the NIH BRAIN Initiative (NS107616) to R.C.F.

## Author Contributions

All authors designed the studies and wrote the paper. S.C. Song performed experiments and analyzed the data. B. Shen and K. Louie performed the modeling.

## STAR Methods

### LEAD CONTACT AND MATERIALS AVAILABILITY

Further information and requests for resources and reagents should be directed to and will be fulfilled by the Lead Contact, Dr. Robert C. Froemke.

#### Materials Availability Statement

This study did not generate new unique reagents.

### EXPERIMENTAL MODEL AND SUBJECT DETAILS

All procedures were approved under NYU School of Medicine IACUC protocols, in accordance with NIH guidelines. Animals were housed in fully-equipped facilities in either the NYU School of Medicine Skirball Institute or Science Building (New York City). The facilities were operated by the NYU Division of Comparative Medicine. C57BL/6 mice (Jackson Labs; Stock No. 000664) of both sexes were used in all experiments; animals were between postnatal day (P) 26 to P226. Mice were either wildtype C56BL/6J or Cre expressing mice directed with specific promoters (Ssttm2.1(cre)Zjh/J, B6.129P2-Pvalbtm1cre(Arbr)/J, Viptm1(cre)Zjh/J, Ndnftm1.1(cre)Rudy/J, B6;C3-Tg(Scnn1a-Cre)3Aibs/J; or B6(Cg)-Cux2tm3.1(cre/ERT2)Mull/Mmmh from the Mutant Mouse Resource & Research Center) to target specific brain regions or neuron types and were either inbred or crossed with B6.Cg-Gt(ROSA)26Sortm32(CAG-Cop4*H134R/EYFP)Hze/J to express channelrhodopsin co-tagged with EYFP (ChR2-EYFP).

### METHOD DETAILS

#### Viral Injections

For some animals, stereotaxic injections of adeno-associated virus (AAV) encoding AAV1.hSyn.hChR2(H134R).EYFP(Addgene) or AAV1.EF1α.FLOX.ChR2.EYFP(Addgene) (ChR2-EYPF) were necessary. Animals were anesthetized with 1.5-2% isoflurane and head fixed on a stereotaxic to make targeted injections into primary auditory cortex (A/P: −2.54, M/L: 4.5, D/V: −0.3) (coordinates are distance, in mm, from bregma and surface of the brain). Injections were made with a Nanoject III (Drummond Scientific) and a freshly pulled micropipette to a ~20 μm diameter. Animals were allowed to recover and virus allowed to express for at least 2 weeks prior to sacrifice and experimentation.

#### Slice Preparation

Acute brain slices were prepared from mice after deep anesthetization with 5% isoflurane. Mice were cardially perfused with ice cold, oxygenated (95% O2/ 5% CO2), sucrose cutting buffer containing (in mM): 87 NaCl, 75 sucrose, 2.5 KCl, 1.25 NaH2PO4, 0.5 CaCl2, 25 NaHCO3, 1.3 ascorbic acid and 10 D-Glucose. After quick decapitation and brain extraction, coronal slices (250 μm thick) were made on a vibrating blade microtome (Leica VT1200s) while submerged in ice cold, oxygenated, sucrose cutting buffer. Slices were then transferred to an incubation chamber containing artificial cerebral spinal fluid (ACSF) consisting of (in mM): 124 NaCl, 2.5 KCl, 1.5 MgSO4, 1.25 NaH2PO4, 2.5 CaCl2 and 26 NaHCO3. The chamber was heated (~35°C) and oxygenated while slices incubated for ~30 min before allowing to adjust to room temperature for 30+ minutes.

#### Whole-Cell Recordings

For recordings, slices were transferred to a holding chamber that was superfused (2-3 mL/min) with oxygenated and heated ACSF. Patch pipettes were made with borosilicate glass pulled to a resistance of 5.5-7.5 MΩ on a Flaming/Brown P-1000 micropipette puller (Sutter Instruments) and filled with (in mM): 127 K-gluconate, 8 KCl, 10 phosphocreatine, 10 HEPES, 4 Mg-ATP, 0.3 Na-GTP. Recordings were made using a Multiclamp 200B amplifier (Molecular Devices), filtered at 2 kHz, digitized at 10 kHz, and acquired with Clampex 10.7 (Molecular Devices). Whole cell recordings were made from layer 2/3 pyramidal neurons. A bipolar stimulating electrode pulled from theta glass was placed in superficial layer 2, proximal to the recorded neuron. Electrical stimulus currents were generated by an A365 stimulus isolator (WPI) triggered by a TTL pulse from the acquisition software. Optical stimulation was done with a PLS series 5500K Cool White LED (Mightex), projected through the objective, and filtered (Semrock) to ChR2 activation wavelength. Data were analyzed using custom written code on Matlab (Mathworks) or Prism (GraphPad).

The induction protocol for STDP was similar to previous studies performed from our lab (D’amour and Froemke, 2015; Field et al., 2020). Baseline currents were measured in voltage-clamp mode for 5+ minutes at a rate of once per minute. For induction, recordings were switched to current-clamp mode to allow the neuron to spike. Utilizing the stimulating electrode, we triggered afferents to layer 2/3 pyramidal neurons and paired presynaptic input with brief postsynaptic depolarization to drive a single action potential in current clamp (**Figure 1B**). The interval between presynaptic stimulation and the postsynaptic current injection was 5 ms. Current amplitude was adjusted for each neuron to reliable trigger an action potential (500 pA to 2.5 nA for a duration of 5 ms). The presynaptic stimulation and postsynaptic pairing was repeated once every 2 seconds for a total of 100 times before returning to voltage-clamp mode and recording ISPCs for up to 30 minutes at a rate of once per minute. Cells had to maintain stable access and input resistances (<30% change from baseline) to be included in final analysis.

#### Numerical Simulations

We tested the function of iLTP in a decision circuit modified from Shen et al. (2022), consisting of different types of units (*R, G*, and *D*) with each unit respectively modelling cortical pyramidal neurons (*R*), SST+/PV+ interneurons (*G*), and VIP interneurons (*D*). The model predicts the temporal dynamic of neural firing rates by using three sets of differential equations. Connection weights in the differential equations are adjusted to model the impact of STDP, including eLTP on the excitatory projections from E to E and the iLTP on the inhibitory connection from I to E:

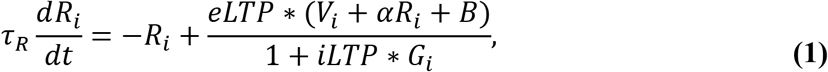

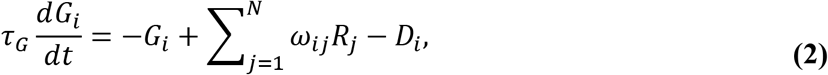

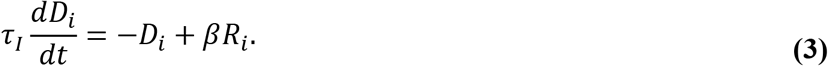

where i = 1, 2 designates the 2 choice alternatives, each of which receive selective input *V_i_* and non-selective baseline input *B. τ_R_*, *τ_G_*, and *τ_D_* are the time constants for the *R*, *G*, and *D* units, which were set to the same value (100 ms). The weights *ω_ij_* represent the coupling strength between excitatory units *R_j_* and gain control units *G*_i_, with its value set as 1.00; the parameter *α* controls the strength of recurrent self-excitation on *R* units, and was set as 15.00. Finally, *β* weights the coupling strength between the excitatory *R_i_* and the disinhibitory *D_i_* units and is presumed to be under external (task-triggered) control. During representation, *β* was set as zero; during choice, *β* was set as 1.00.

The Runge-Kutta method in MATLAB (MathWorks) was used to numerically implement the differential equations at time step of 1 ms. Decision threshold for choice was fixed at 70 Hz. The input values were given based on the percentage of difference of the input stimuli (*c’*), where *V_1_* = *S*\*(1 + *c’*) and *V_2_* = *S*\*(1 – *c’*). *S* indicates the scale of the input and was set as 576; *c’* can be also noted as *c’* = (|V1-V2|) / (V1+V2))). *c’* was set as 6.4% in the visualization of representation dynamics and distributions (**Figure 3C,D** and the first column of **Figure 4**). In the testing of choice behavior (**Figure 3F** and the second and the third columns of **Figure 4**), *c’* varies from small to large differences ([.2, .4, .8, 1.6, 3.2, 6.4, 12.8, 25.6, 51.2]%); *c’* was set as 3.2% in the testing of input noise levels in the fourth column of **Figure 4**. The baseline non-selective input (*B*) was set as zero. For implementation of noise, two independent chains of additive random noise were added onto *V_1_* and *V_2_*, with each chain drawn from a Gaussian distribution with zero mean and standard deviation *S*σ_Input_* at every 5 ms. *σ_Input_* was set as 1/3 in the visualization of neural dynamics and representational distributions (**Figure 3C-E** and the first column of **Figure 4**). *σ_Input_* was tuned to be .75 in the visualization of choice behavior (**Figure 3F** and the second and the third columns of **Figure 4**). *σ_Input_* varies from 0 to 2 at a step of .02 in the testing of different levels of input noise (the fourth column of **Figure 4**). In addition to the noise on input, we preserved a small amount of intrinsic stochasticity in the circuit but not to overwrite the effect of input noise. We kept the endogenous noise that was added on to the neural activities as low as *σ* = .1 (Shen et al., 2022). This noise source is different from the source of input noise *σ_Input_*. Two levels of LTP (iLTP or eLTP = 1.00 and 2.00) were picked to study the neural representation shown in the first column of **Figure 4**. Five levels of LTP (1.00, 1.75, 2.50, 3.25, 4.00) are compared in the second and the third columns of **Figure 4**. Finally, two levels of LTP (1.00 and 4.00) were picked to show the choice accuracy over different levels of input noise in the fourth column of **Figure 4**.

### QUANTIFICATION AND STATISTICAL ANALYSIS

Student’s t-test was used for comparisons between IPSC amplitudes before vs after spike pairing, with paired or unpaired tests used when appropriate. Statistical analyses were performed using Prism 6.0 GraphPad and MATLAB (MathWorks). Statistical tests used, p-values, and the number of cells are reported in the main text describing each figure. All quantifications are the result of data from at least 3 different animals, unless otherwise indicated. Data reported in the text are generally shown as mean ± standard error of the mean (s.e.m), unless otherwise indicated.

### DATA AND CODE AVAILABILITY

Upon request to the Lead Contact, data are immediately available.

